# Lateral lipid packing governs bilayer solubilization by styrene-maleic acid copolymers: a case study with cardiolipin-containing membranes

**DOI:** 10.1101/2025.06.23.659572

**Authors:** Joseph C. Iovine, Nathan N. Alder

**Author notes:** Corresponding Author: Dr. Nathan N. Alder, 91 N. Eagleville Rd, Storrs, CT 06269-3125.

## Abstract

Styrene-maleic acid (SMA) copolymers are powerful tools for the detergent-free solubilization of biological membranes. Yet the influence of specific lipids on SMA activity remains an open question. Here, we examined the effects of the mitochondriaspecific phospholipid cardiolipin on SMA-mediated membrane solubilization and their ability to form SMA-bound nanodiscs. To this end, we prepared a series of model membranes with cardiolipin and other test lipids with comparable surface charge and lateral packing characteristics. Using multiple independent experimental approaches, we found that cardiolipin inhibited SMA solubilization. Our results indicate that this effect was not attributable to headgroup charge effects, but rather to cardiolipininduced increase in lateral packing pressure at the interfacial region. Reduction of this lateral packing pressure using bilayer-active alcohols partially restored SMA solubilization. Our results highlight the importance of lipid geometry and packing in SMA nanodisc formation and could help guide the design of copolymers tailored to specific membranes.

## 1. Introduction

At the forefront of membrane biology research are synthetic membrane-interactive amphipathic polymers that solubilize membranes into soluble nanoscale discs (1, 2). The development of this class of compounds was initiated with styrene-maleic acid (SMA) copolymers (3, 4), and to this day SMA is the predominant copolymer being used in basic and applied studies (5, 6). The amphipathic SMA polymer contains hydrophobic (styrene) and hydrophilic (maleic acid) pendant groups, typically in molar ratios of 3:1 or 2:1. SMA partitions into membranes, forming nanoscale, discoidal lipoprotein particles (nanodiscs) stabilized at the rim by the polymer belt in which the styrene moieties interact with the nonpolar lipid acyl chains and the maleic acid groups interact with the aqueous environment (4, 7–9). SMA effectively forms nanodiscs from pure lipid model membranes, from proteoliposomes, and from native cellular membranes (10–12). The resulting nanoparticles are termed ‘native nanodiscs’ or SMA-lipoprotein particles (SMALPs). From a practical standpoint, amphipathic polymers like SMA allow for the one-step, detergent-free extraction of membrane lipids and proteins into soluble, experimentally tractable systems that can be used to study membrane proteins in their native lipid environments (13–24) and can be used as nanocarriers for drug compounds that enhance bioavailability and allow controlled release (25–28).

To understand the mechanism by which SMA copolymers form nanodiscs, many studies have used carefully defined experimental conditions to measure the kinetics and yield of SMALP formation. For example, SMA activity is strongly modulated by solution conditions, including pH, ionic strength, and concentration of divalent cations (29–31). Another key factor is the way in which the physical and chemical properties of the target membrane may influence SMA efficiency. This can be tested using reductionist, lipid-only model membranes (typically liposomes) with defined lipid composition in which the headgroup and acyl chain features are controlled. Based on studies using this approach (8, 32–35), the following general patterns emerge: (1) SMA solubilization rate is greater when membranes are in the in the liquid crystalline phase (T>T_M_) than in the gel phase (T<T_M_); (2) Solubilization rate generally increases with temperature; (3) Solubilization rate decreases with increasing acyl chain length (greater membrane thickness); (4) Unscreened negative surface charge of lipid headgroups inhibits SMA solubilization due to the electrostatic repulsion with the anionic maleic acid pendant groups; and (5) Increased lateral packing pressure in the membrane nonpolar domain, caused by acyl chain unsaturation and/or lipids with an inverted conical (H_II_) molecular geometry, inhibit SMA insertion into the bilayer. Notably, when SMA is used to solubilize model or natural membranes, the lipid composition of the resulting SMALPs generally reflects that of the originating membrane (8, 13, 14, 16, 32), suggesting that it is the collective physical properties of the membrane that determine SMA efficiency, not individual lipid constituents.

An important question that remains in SMA research is the efficiency of this copolymer in solubilizing different types of cellular membranes. SMA shows different activity toward plasma and organellar membranes (36–39), which may be influenced in part by the presence of specific lipids. Indeed, understanding how membrane properties influence SMA activity is critical for the design of copolymer variants that are better able to solubilize different membranes. In the present study, we addressed the ability of 3:1 SMA to solubilize membranes containing cardiolipin (CL), the signature lipid of mitochondria, which is enriched in the mitochondrial inner membrane (MIM) (40). The lipid content of the MIM is almost exclusively glycerophospholipids, which predominantly include the bilayer-forming lipid phosphatidylcholine (PC) and non-bilayer lipids phosphatidylethanolamine (PE) and CL (41, 42). These non-bilayer lipids have an inverted conical molecular geometry that allows them to accumulate in and stabilize regions of negative curvature (concave toward the headgroup), which is essential for maintaining the highly curved structures of the MIM (43–46). In addition to regulating membrane morphology, CL assumes a particularly critical role in the MIM, where it supports the assembly and function of protein complexes and serves as a signaling molecule for processes like mitophagy and apoptosis (47–52).

CL is a phospholipid with an unusual dimer-like structure, containing a two phosphate diester headgroup and a tetra-acyl nonpolar region of four acyl chains that are highly unsaturated (53). As such, it contains two key features – a negatively-charged headgroup and inverted conical geometry – that are likely to inhibit SMA activity. In a recent study, Janson et al. evaluated the ability of amphipathic copolymers to solubilize MIM-mimetic membranes (54). Using model membranes made of naturally sourced lipids, they found that CL inhibited SMA-mediated solubilization and found that the underlying basis for this inhibition was the anionic charge repulsion between polymer and lipid. Here we take a different tack, using model membranes of synthetic lipids with uniform acyl composition. While we also find that CL inhibits SMALP formation, our experimental system points to a dominant role of lateral packing pressure, not charge repulsion, in this inhibition. This study offers key insights into the resistance of CL-containing bilayers to SMA solubilization and, compared with previous work, suggests how variations in SMA and CL acyl composition might influence this resistance.

## 2. Materials and Methods

### 2.1 Reagents and Phospholipids

Lipids chosen for this study were synthetic, monounsaturated phospholipids obtained from Avanti Polar Lipids (Alabaster, AL, USA) as chloroform stocks. These included 16:0-18:1 POPC (1-palmitoyl-2-oleoyl-glycero-3-phosphocholine), 16:0-18:1 POPE (1-palmitoyl-2-oleoyl-sn-glycero-3-phosphoethanolamine), 16:0-18:1 POPG (1-palmitoyl-2-oleoyl-sn-glycero-3-phospho-(1’-rac-glycerol) (sodium salt)), fully monounsaturated 18:1 CL (TOCL, 1’,3’-bis[1,2-dioleoyl-sn-glycero-3-phospho]-glycerol (sodium salt)), and 18:1 monolysocardiolipin (MLCL). All lipid stocks were stored at −20°C in clear glass vials with Teflon-lined cap closures.

Styrene-maleic acid anhydride 3000 was obtained from Cray Valley, LLC and converted into styrene-maleic acid copolymer (3:1 SMA) using the sodium hydroxide hydrolysis method as described (55). The fluorescent probe 6-Dodecanoyl-2-Dimethylaminonaphthalene (laurdan) was purchased from ThermoFisher. The fluorescent probes N,N-Dimethyl-6-propionyl-2-naphthylamine (prodan) and 8-Hydroxypyrene-1,3,6-trisulfonic acid trisodium salt (HPTS) were purchased from Millipore-Sigma at purity grades of ≥98.0% and ≥96.0%, respectively. All other chemicals were reagent grade, obtained from Millipore-Sigma. All solutions were prepared with ultrapure water (Millipore Advantage A10 system; resistivity 18.2 MΩ•cm @ 25°C; total oxidizable carbon ≤ 4 ppb).

### 2.2. Preparation of liposomes

Large unilamellar vesicles (LUVs) were prepared by thin film rehydration of lipid mixtures and extrusion as described (56). Chloroform lipid stocks were mixed at the appropriate molar ratios in thoroughly cleaned and dried high-strength glass test tubes. The chloroform was evaporated under a gentle nitrogen stream and the resulting lipid film was dried in a vacuum chamber (Eppendorf vacufuge) for a minimum of two hours. Lipid films were then hydrated in lipid buffer (300 mM NaCl, 20 mM HEPES, pH 7.5) by gentle swirling to form multilamellar lipid vesicles (MLVs). LUVs were then prepared by subjecting MLVs to five freeze-thaw cycles (snap-freezing in liquid nitrogen and thawing in a 45°C water bath) followed by extrusion (passage through a polycarbonate filter exactly 23 times) using a mini extruder (Avanti). LUVs prepared for particle size assays and laurdan/prodan measurements were extruded using 0.1 μm filters; those used for HPTS measurements were extruded using 0.8 μm filters. Larger vesicles were used for HPTS assays to maximize intravesicular volume (and therefore the total amount of probe per vesicle) to improve the signal-to-noise fluorescent readout. This process resulted in single monodisperse populations of LUVs, confirmed by dynamic light scattering measurements (below). The total concentration of lipids recovered from the extrusion process was determined by the colorimetric Stewart assay using an ammonium ferrothiocyanate assay as described (57) against lipid standards based on the mole fraction of lipid types in each preparation.

### 2.3. Dynamic light scattering and zeta potential measurements

All dynamic light scattering (DLS) and zeta potential (ζ) measurements were done with a Zetasizer Pro instrument (Malvern Panalytical). Each independent measurement consisted of freshly prepared LUVs, analyzed in triplicate (three technical replicates). DLS measurements were made in low volume quartz cuvettes (Malvern model DTS2112) with LUVs diluted to a final total lipid concentration of 800 μM in lipid buffer equilibrated to 25°C prior to measurement. Data analysis was done using software associated with the instrument (ZS XPLORER version 2.0.0.98). Z-averages were calculated from the autocorrelation function for each technical replicate, and values were averaged across replicates for each independent trial. For particle size assays evaluating the SMA-mediated membrane solubilization, LUVs were measured for size distribution by DLS before and after SMA incubation. For these experiments, SMA was added to LUVs at a final SMA:lipid ratio of 1:2.92 (mol:mol) and incubated at room temperature for 15 minutes. ζ measurements were performed using disposable folded capillary cells (Malvern model DTS1070, polycarbonate with gold-plated copper electrodes). For these experiments, LUVs were diluted to a final total lipid concentration of 200 μM in 30 mM NaCl, 20 mM HEPES, pH 7.5.

### 2.4. Fluorescence spectroscopy

All steady-state fluorescence measurements were done with a Fluorolog QM-75-22-C photon counting spectrofluorometer (HORIBA Scientific). For all fluorescence experiments, each treatment group was measured a minimum of n=3 times with freshly prepared LUV samples for each replicate.

For experiments using laurdan and prodan probes, fluorescent LUV samples were prepared as described (58–64) by adding solutions of probe in concentrated stocks to pre-formed vesicles. LUVs were diluted to a total lipid concentration of 50 μM in lipid buffer, incubated with 1 μM laurdan or prodan (added from a 1 mM DMSO stock) for a minimum of 1h with gentle rotation and protection from light. For spectral measurements, samples were loaded into a 500 μl quartz microcuvette (Starna Cells) and emission spectra were recorded (λ_ex_ =350 nm; λ_em_ = 370-600 nm). All spectra of fluorescent samples were corrected for background by scalar subtraction of scans conducted with non-fluorescent LUVs. The Generalized Polarization (GP) parameter for laurdan- and prodan-containing samples was calculated using the equation *GP* = (*I*_440_ − *I*_490_)⁄(*I*_440_ + *I*_490_) as described (65, 66).

For time course measurements of membrane permeabilization, LUVs with the HPTS probe loaded into the lumen were prepared by freeze-thaw as described (67, 68). LUVs at a total lipid concentration of 4 mM in buffer (300 mM NaCl, 20 mM HEPES, pH 7.5) were incubated with 6 mM HPTS (added from a 100 mM stock in the same buffer used for LUVs) and subject to three freeze-thaw cycles (snap-freezing in liquid nitrogen and thawing in a 25°C water bath for 5 min). HPTS-loaded LUVs were then passed through a PD-10 desalting column (Cytiva) to remove all free dye, diluted to a total lipid concentration of 1 mM in lipid buffer for storage. For spectral measurements, HPTS-loaded LUVs were diluted to a total lipid concentration of 100 μM in 3.5 ml lipid buffer (adjusted to pH 8.5) and placed in a 4 ml cuvette (Starna Cells) equipped with a spin disc. Time course measurements on stirred samples were made using the synchronous function (emission measurement at λ_em_ = 510 nm with alternating excitation wavelengths of λ_ex_ = 405 nm and λ_ex_ = 458 nm). Measurements were repeated a total of 60 times for each run for a total experiment runtime of approximately 5 min 40 sec. After ten repeats, SMA was added to a final SMA:lipid ratio of 1:2.92 (mol:mol). Prior to SMA addition, an HPTS emission spectrum was obtained for each sample to ensure that the luminal pH remained consistent with the pH 7.5 lipid buffer used for LUV preparation (i.e., that vesicles retained the pH gradient). For each time course reading, the excitation ratio corresponding to *I*_405_/*I*_458_ was converted into pH values based on an independently prepared pH standard curve using a second order polynomial fit.

## 3. Results and Discussion

### 3.1. Characterization of lipid vesicles: size and surface charge

To evaluate the effects of CL on SMA-dependent membrane solubilization, we prepared a series of LUVs composed of synthetic glycerophospholipids (**Table 1**). The lipid compositions were designed to measure the effects of CL as well as other test lipids that have key features of CL. The first feature is headgroup charge. Glycerophospholipids have headgroups that are either charge-neutral (zwitterionic) or bear a net negative charge from the phosphate and other functional groups. Most negatively charged phospholipids are monoanionic (formal charge −1); however, CL, containing a two-phosphate headgroup, is dianionic (formal charge −2) over a wide pH range, as has been confirmed using model membrane systems (69–72). The second feature is molecular geometry. Lipid shape can be described by the size of the polar headgroup relative to the nonpolar hydrocarbon tails, quantifiable by the dimensionless molecular packing parameter, *S* = *v* / *a*_o_*l*_c_, where *v* is the volume occupancy of the acyl tails, *a*_o_ is the optimal cross-sectional area of the headgroup, and *l*_c_ is the critical (maximally extended) length of the acyl tails (73). Lipids with comparable headgroup and hydrocarbon tail sizes assume a cylindrical geometry (S ∼ 1) that favors bilayer formation, whereas those with a headgroup that is smaller than the time-averaged volume of the hydrocarbon tails assume an inverted conical geometry (S > 1) that favors nonlamellar, inverted hexagonal (H_II_) phases (74). CL, containing four acyl tails that are typically unsaturated, has a large nonpolar volume relative to its headgroup and is therefore considered an inverted conical lipid (75–77).

**Table 1.** Phospholipid species used in this study and their relevant properties.

| Variable Lipid | Formal Headgroup Charge | Molecular Geometry | LUV Composition (mol % POPC/mol % Variable Lipid) | Hydrodynamic Size (nm)<sup>a,b</sup> | Zeta Potential (mV)<sup>b</sup> |
| --- | --- | --- | --- | --- | --- |
| POPC | 0 | Cylindrical | 100/0 | 148 (1.7) | -4.97 (0.09) |
| TOCL | -2 | Cylindrical / Inverse Conical | 90/10 | 134 (10) | -38.2 (2.35) |
|  |  |  | 80/20 | 137 (8.2) | -45.7 (1.55) |
|  |  |  | 70/30 | 136 (13) | -48.5 (0.09) |
| POPG | -1 | Cylindrical | 90/10 | 144 (4.2) | -31.2 (1.52) |
|  |  |  | 80/20 | 139 (5.6) | -37.7 (0.93) |
|  |  |  | 70/30 | 136 (2.7) | -42.7 (1.57) |
| POPE | 0 | Inverse Conical | 90/10 | 159 (4.3) | -4.88 (0.95) |
|  |  |  | 80/20 | 159 (7.1) | -4.87 (0.86) |
|  |  |  | 70/30 | 163 (3.7) | -6.78 (0.25) |
| MLCL | -2 | Cylindrical / Conical | 90/10 | 135 (16) | -35.5 (0.53) |
|  |  |  | 80/20 | 137 (11) | -43.6 (1.42) |
|  |  |  | 70/30 | 133 (5.7) | -45.7 (2.79) |

<sup>a</sup>Hydrodynamic sizes were measured for all LUVs in particle size and Laurdan/Prodan assays
<sup>b</sup>Errors indicate standard deviations

Our reference LUVs were composed of 100% POPC (16:0/18:1 PC), which served as a suitable control for analyzing the effects of variable test lipids because PC is the predominant phospholipid of eukaryotic membranes (42) and vesicles made of PC are efficiently solubilized by SMA (8, 32–35). The effects of variable lipids were evaluated by preparing LUVs with 10, 20, and 30 mol% of each test lipid in a POPC host background. Our first test lipid was TOCL (18:1 CL), which served as a model cardiolipin variant because its headgroup is dianionic in the physiological pH range (69–72) and its four monounsaturated acyl chains impart H_II_ propensity (78). Our second test lipid was POPG (16:0/18:1 PG), a bilayer-forming lipid chosen because its monoanionic headgroup imparts a net negative surface charge to the bilayer. Our third test lipid was POPE (16:0/18:1 PE), a zwitterionic lipid chosen for its inverted conical (H_II_-preferring) geometry that causes increased packing pressure in the acyl region. Our fourth test lipid was 18:1 MLCL, a tri-acyl cardiolipin with a headgroup structurally identical to TOCL and with the same ionization properties (71) but lacking one 18:1 acyl chain at the *sn*-2 position so it assumes a more cylindrical geometry (79). Note that for this study, we chose to use synthetic lipids instead of naturally sourced ones because of their purity, their defined acyl composition, and their lot-to-lot consistency.

Each LUV type containing 10-30 mol% test lipids was characterized for key parameters. First, we measured the hydrodynamic diameters of LUVs by dynamic light scattering (DLS) (**Table 1**). All LUV preparations had a monodisperse size distribution and reasonably low polydispersity index (**Supplementary Table 1**), indicating uniform vesicle sizes. The measured LUV diameters were all larger than the 100 nm nominal pore size used for extrusion, which is typical for LUVs prepared by this method (80). This could result from many factors, including the fact that DLS may overestimate vesicle diameter by accounting for bound water layers, LUVs can deform during extrusion and relax to a larger size, and bilayer-preferring lipids (S ∼ 1) or H_II_-preferring lipids (S > 1) on the outer leaflet could favor lower curvature, larger vesicle sizes.

Next, we characterized surface charge by zeta potential (ζ) measurements (**Table 1**). Notably, our LUVs consisting of purely electroneutral lipid (POPC and POPE) showed slightly negative ζ values. This modest negative surface charge may reflect asymmetric solvent exposure of the phosphate versus choline or ethanolamine groups, or weak preferential adsorption of anions versus cations – phenomena that have been reported in previous studies (81–84). Compared with our POPC host lipids, our LUVs with test lipids revealed the following expected patterns: (i) LUVs with the zwitterionic test lipid POPE had a ζ near zero, (ii) LUVs containing anionic test lipids POPG, TOCL, and MLCL had ζ values that became progressively more negative with increasing anionic lipid concentration, and (iii) LUVs with dianionic test lipids (TOCL and MLCL) had more negative ζ values than those with monoanionic test lipids (POPG). Our measured ζ values are consistent with published values for LUVs from our lab (71) and others (85).

### 3.2. Characterization of lipid vesicles: lateral packing properties

We next addressed how our different LUV preparations differed with respect to lateral lipid packing at different membrane depths using the microenvironment-sensitive probes laurdan and prodan. Both reporters contain the same naphthalene-based solvatochromic fluorescent moiety. However, prodan contains a short propionyl tail and resides superficially in the bilayer, near the level of phospholipid glycerols and bulk solvent, whereas laurdan contains a longer lauric tail, residing deeper in the bilayer near the polar-apolar boundary (86, 87). The emission spectra of laurdan and prodan are sensitive to lipid packing, based on the interfacial exposure of the naphthalene moiety to water molecules with resulting dipolar relaxation of the probe. Lateral packing is quantified by the intensities of two emission peaks, one toward the blue end of the spectrum (λ_em_=440 nm) that increases in liquid ordered (*l*_o_) phases (less probe hydration), and one toward the green end of the spectrum (λ_em_=490 nm) that increases in liquid disordered (*l*_d_) phases (more probe hydration). The relative intensities of these peaks are quantified as the generalized polarization (GP) function (see Section 2.4), which can assume limiting values from −1 to +1, with more negative and more positive values corresponding to more disordered (*l*_d_-like) and ordered (*l*_o_-like), respectively (88). Importantly, all lipids used in this study have at least one unsaturated acyl chain and therefore have main transition melting temperatures (T_m_) below 0°C. All LUVs will therefore be in the *l*_d_ phase under the conditions tested; however, given that prodan and laurdan show continuous emission red shifts with increasing temperature in the *l*_d_ phase (89), any lipid-specific changes in lateral packing should still be detectable, even though all bilayers are in the liquid-crystalline phase. Furthermore, prodan and laurdan spectral shifts are independent of headgroup chemical composition or formal charge (88); therefore, their GP values specifically reflect phase state and lateral packing.

In our protocol, we sought to measure only the solvent-exposed monolayer of the LUVs with these probes, reasoning that this would be the most relevant leaflet to analyze for SMA membrane engagement. We therefore prepared prodan- and laurdan-labeled LUVs by the addition of probe stocks to pre-formed vesicles, a method that has been thoroughly validated (58–64). The majority of probes that partition into the outer leaflet will remain in that leaflet given that transbilayer diffusion is highly unlikely, and this approach provides fluorescent signal only from membrane-bound probes because prodan and laurdan emission is negligible in aqueous media.

The measured prodan and laurdan GP values for LUVs containing the different lipid blends used in our study are shown in **Figure 1**. In general, the prodan GP values for these PC-based LUVs are lower than the corresponding laurdan GP values, reflecting the fact that prodan is in a more polar environment near the bulk aqueous phase, subject to the dipolar relaxation of more freely rotating water dipoles (89). Using our POPC-only LUVs as a comparator, our measured prodan GP values (∼ −0.3) and laurdan GP values (∼ −0.12) are in good agreement with previously reported values for PC-containing vesicles (90–92).

**Figure 1.**
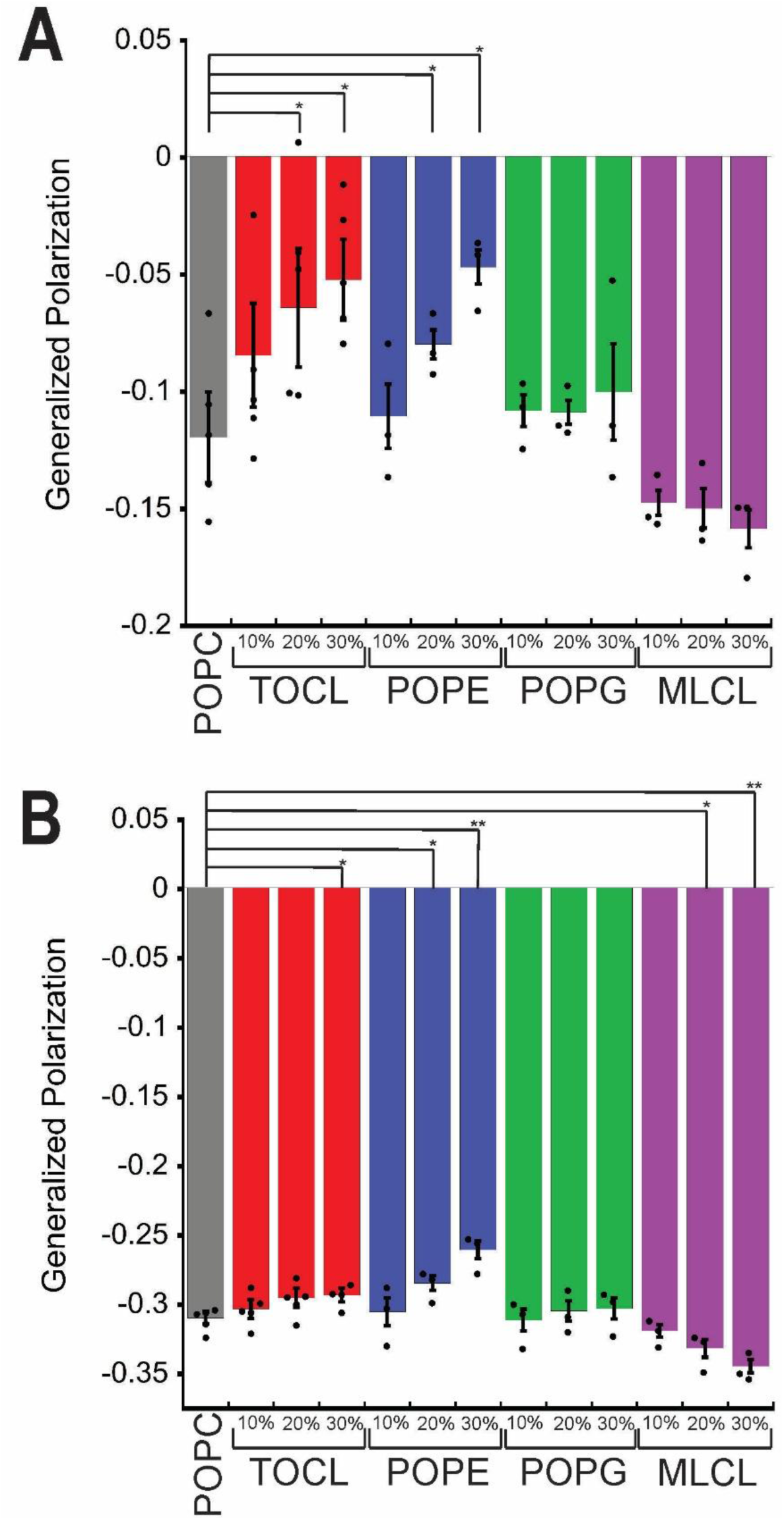
GP values of LUVs with varying lipid composition. GP values were determined using (A) laurdan and (B) prodan fluorescence. LUVs were composed of POPC alone (gray) or with different mol% of test lipids: TOCL (red), POPE (blue), POPG (green), or MLCL (violet). Individual data points (circles) represent experimental replicates (n = 3-5) and bars represent the mean ± SD. Statistical comparisons were made to POPC-only controls using unpaired t-tests (*P < 0.05).

As expected, given the similar cylindrical geometry of POPG and POPC, the former had no detectable effect on local hydration in the polar headgroup region (prodan, **Figure 1A**) or near the polar-apolar boundary (laurdan, **Figure 1B**). The most prominent effect of our test lipid series was that inverted conical lipids (TOCL and POPE) increased lipid packing, to varying degrees, at different depths of the interface. Empirical and computational analyses of lateral lipid pressure profiles show that H_II_-preferring lipids in bilayers generally increase the positive pressure in the acyl chain region while decreasing pressure at the level of the headgroups, given the small cross-sectional area of the headgroups relative to the wider volume occupied by lipid tails (93–96). In our LUVs, increasing POPE content increased GP values of both prodan and laurdan, consistent with previous studies using these probes with PE-containing membranes (90, 97). This may be attributed to PE species inducing more ordered lipid packing near the membrane surface, due in part to the ability of the primary amine-containing headgroup to act as a hydrogen bond donor that promotes noncovalent interactions with neighboring lipids and water molecules (98, 99). By comparison, we found that TOCL only modestly increased order in the headgroup region, which may differ from POPE because the only hydrogen bond donor of CL, the secondary hydroxyl of the central glycerol, is conformationally restricted and not likely to stabilize headgroup interactions as strongly as the PE headgroup (77). Yet TOCL significantly increased order at the polar-apolar boundary, consistent with previous reports that CL increases laurdan GP values in bilayers, likely correlated with its inverted conical geometry that produces positive pressures below the level of the headgroups (91, 100–102). Interestingly, by contrast, for MLCL-containing membranes, the prodan GP value was significantly lower than that of the POPC-only control and the laurdan GP value trended lower (though not significantly so) than that of the control, indicating greater disorder at the interface. In MLCL, the replacement of one acyl chain on a headgroup glycerol with a hydroxyl group promotes conformational flexibility and reduces the local steric bulk enough that it approximates a cylindrical lipid, a phenomenon supported by lateral pressure profiles showing reduced magnitude of interfacial tension and positive pressure in the acyl chain region when comparing TOCL and MLCL (79).

### 3.3. Analysis of lipid-dependent SMA solubilization by particle size assays

Having characterized the key parameters of our LUVs with different lipid compositions, we then evaluated the effects of our different test lipids on SMA-dependent membrane solubilization. To this end, we used a particle size assay based on dynamic light scattering (DLS), a noninvasive technique suitable for measuring the SMA-dependent disruption of lipid vesicles to form SMALPs (32, 103). Lipid compositions that are amenable to SMA solubilization will undergo a DLS-detectable reduction from the starting size of our lipid vesicles (LUVs, ∅ ∼ 150 nm) to the expected sizes for SMALPs containing lipids (∅ ∼ 8-12 nm) (3–5, 7, 32) under conditions of saturating SMA concentrations. To illustrate, POPC-only control LUVs have DLS-detected monodisperse size distributions centered at 150 nm for intact LUVs and centered at 17 nm following solubilization by the SMA copolymer (**Figure 2**).

**Figure 2.**
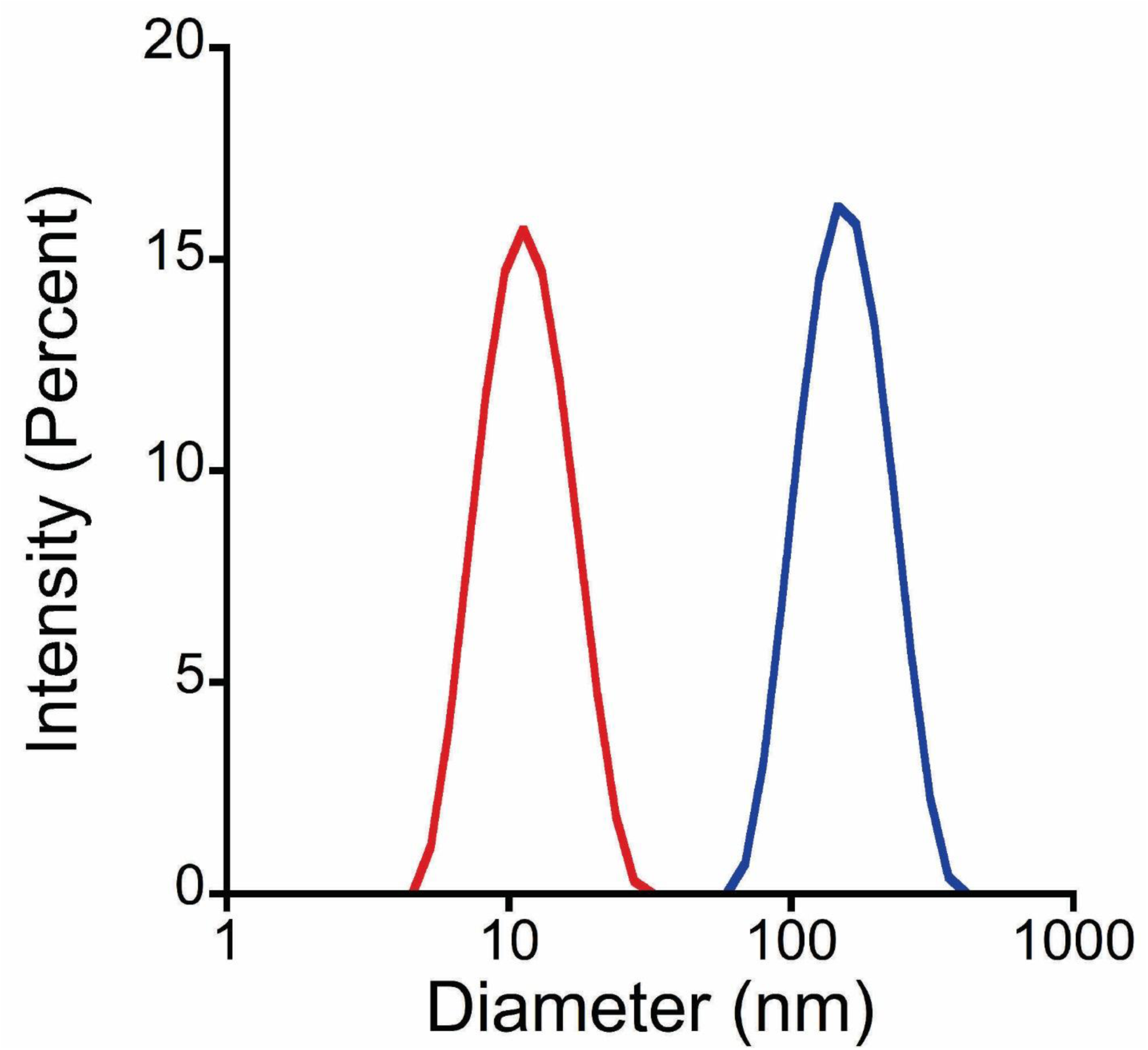
Intensity-weighted DLS size distribution of POPC-only LUVs without SMA addition (blue; Z-average diameter 148 nm; PDI 0.174) and after SMA addition (red; Z-average diameter 17 nm; PDI 0.264).

We performed our SMA solubilization assay using LUVs with a host background of POPC and 0, 10, 20, or 30 mol% test lipid (POPG, POPE, TOCL, or MLCL). Note that for these tests, we used a SMA:lipid molar ratio of 1:2.92 to evaluate SMA solubilization of all LUV lipid compositions. Assuming a molecular weight averaged molar mass (M_w_) of 10 kDa for 3:1 SMA (8), this translates into a SMA:lipid ratio of at least 3:1 (w/w), which is in the range of excess SMA concentration for solubilization of PC-based liposomes (32). For LUVs containing POPC only, or up to 30 mol% POPG, POPE, or MLCL, vesicles were essentially completely converted to SMALPs under these reaction conditions; the only exception was TOCL-containing LUVs, which showed strong inhibition of SMALP formation in a TOCL concentration-dependent manner (**Supplementary Figure S1**). Notably, TOCL-containing samples with inhibited levels of SMALP formation did not form vesicle sizes intermediate between full-size LUVs and SMALPs; rather, they retained bimodal distributions whose relative amounts of SMALP fractions decreased with increasing TOCL. As a single-parameter index of SMA solubilization efficiency, we determined the Z-average, the intensity-weighted average of particles in the sample (**Figure 3**) prior to (blue traces) or after (red traces) incubation with SMA. Whereas LUVs containing test lipids POPE, POPG, and MLCL quantitatively transitioned from full-size vesicles to SMALPs following SMA incubation, those containing TOCL showed inhibition of SMALP formation in a [TOCL]-dependent manner. Hence, among our test lipids, TOCL uniquely inhibited SMA solubilization under these reaction conditions.

**Figure 3.**
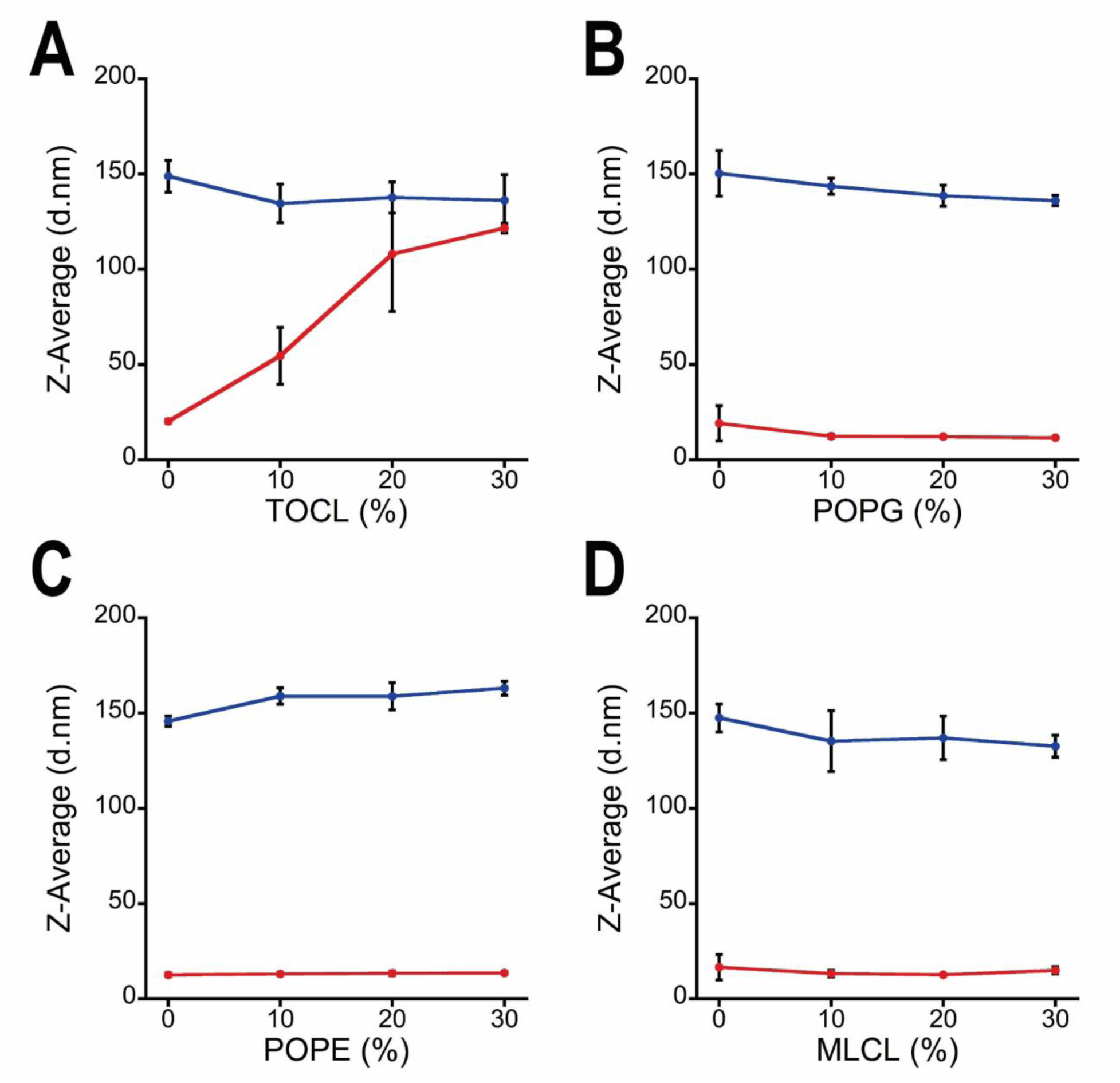
Z-average diameters of LUVs before and after SMA incubation. DLS-measured, intensity-weighted Z-average diameters (mean ± SD, n = 3-5 experimental replicates) for LUVs composed of POPC with the indicated mol% of (A) TOCL, (B) POPG, (C) POPE, and (D) MLCL. Blue lines connect data points prior to SMA treatment; red lines connect values after incubation with SMA (SMA:lipid ratio = 1:2.92 mol:mol, 15 min at RT).

We next addressed whether the inhibitory effect of TOCL was due to suboptimal SMA concentration and/or insufficient time for solubilization (**Figure 4**). To more clearly evaluate the formation of SMALPs, in these tests we quantified the area under the curve (AUC) of the size distribution corresponding to SMALPs in our DLS profiles. First, to test the SMA concentration dependence, we titrated SMA to a concentration up to approximately ten times that used in previous experiments (**Figure 4A**). This titration revealed a pronounced effect of TOCL; specifically, LUVs containing 10 mol% TOCL had a SMA concentration-dependent solubilization profile essentially equal to that of POPC-only controls, whereas LUVs containing 20 and 30 mol% TOCL showed progressive inhibition of SMALP formation over the 15-minute solubilization reaction. Second, to test the time dependence, we compared SMALP formation of the same LUV series with a fixed SMA:lipid ratio (1:2.92 mol:mol) over longer SMA incubation times (up to 60 min) (**Figure 4B**). Even with extended incubation times, LUVs containing 20 mol% and 30 mol% TOCL showed partial and complete inhibition of SMALP formation, respectively. We conclude that in our system TOCL inhibits SMA activity at concentrations at or above 20 mol% in a manner that cannot be overcome with increased [SMA] or extended reaction time. Finally, we determined whether TOCL affected the minimal concentration of SMA required to effect LUV solubilization (**Figure 4C**). Using a high-resolution SMA titration, we found that considerably higher concentrations of SMA were required to fully solubilize LUVs with 20 mol% TOCL relative to POPC controls; however, nanodisc formation began at similar concentrations of SMA regardless of LUV lipid composition (**Figure 4C, inset**). This observation is consistent with a previous study (8) showing that the H_II_ lipid POPE raised the polymer concentration required for complete vesicle solubilization but lowered the threshold for membrane polymer saturation. These results are consistent with a model in which SMA-induced solubilization proceeds via a multistep mechanism, with TOCL interfering primarily with downstream steps – such as membrane remodeling into nanodiscs – with initial membrane binding or polymer saturation likely remaining unaffected.

**Figure 4.**
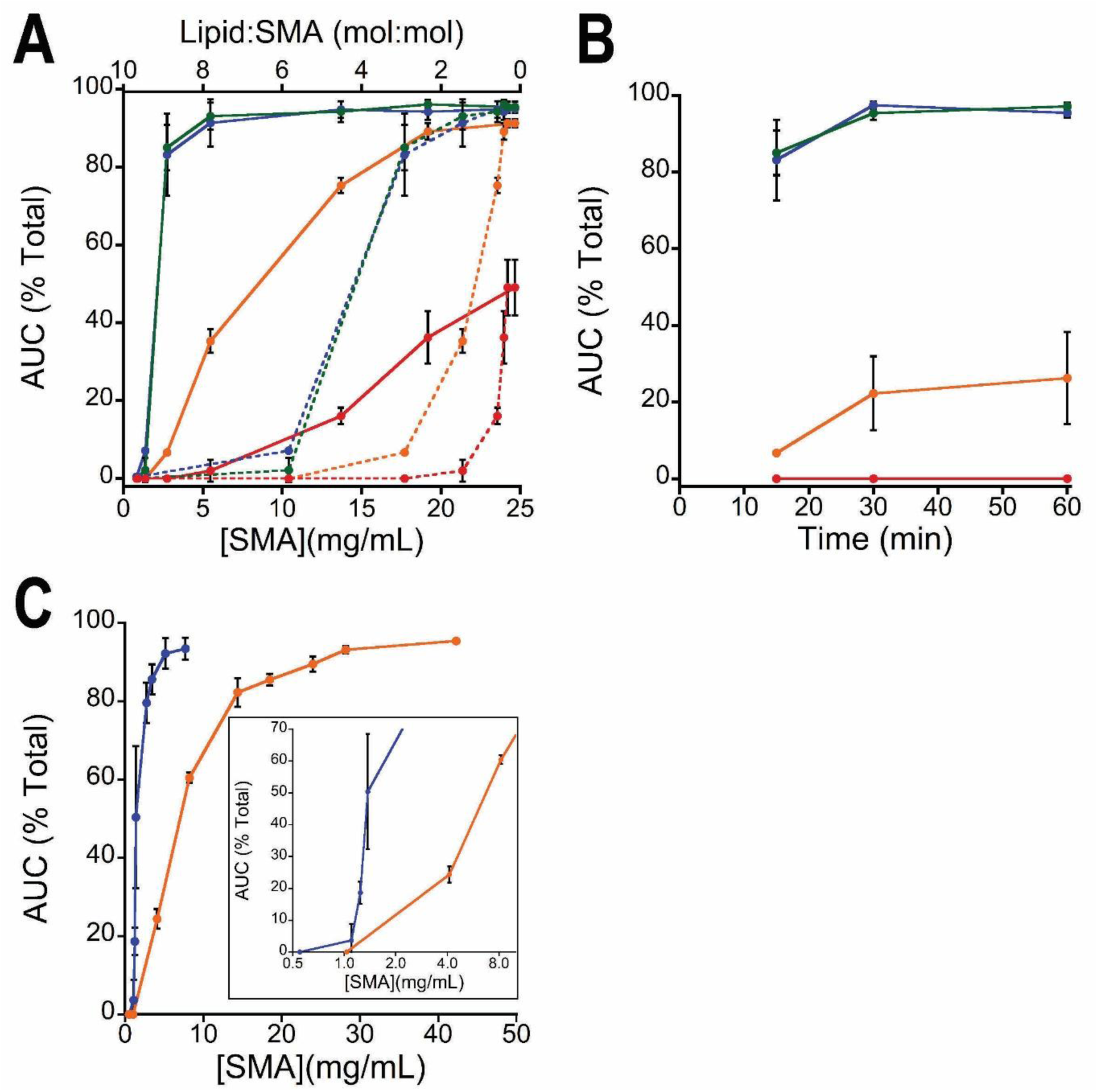
SMA concentration and time dependence of SMALP formation from TOCL-containing vesicles. SMALP formation was assessed using DLS size distribution profiles and quantified as the percentage of total particle area corresponding to the SMALP fraction (∼10 nm diameter), calculated from the area under the curve (AUC) for each sample. LUVs were comprised of POPC only (blue), 10 mol% TOCL (green), 20 mol% TOCL (orange), and 30 mol% (red). Individual data points represent means ± SD (n = 3-5 experimental replicates). (A) Double X plot showing SMA concentration dependence of solubilization, quantified relative to total [SMA] (lower X axis) and the lipid:SMA molar ratio (upper X axis). Vesicles were incubated 15 min at RT with the indicated concentration and SMALP AUC percentages were determined. (B) Time dependence of solubilization. Vesicles were incubated with SMA at a final SMA:lipid ratio of 1:2.92 mol:mol at RT, and SMALP AUC percentages were measured following incubation at the indicated time points. (C) Fine-scale SMA titration showing nanodisc formation at the lowest SMA concentrations (inset). Individual data points represent means ± SD (n = 3-5 experimental replicates).

TOCL has two properties that are known to inhibit key steps of the SMA mechanism (8, 31, 32, 54). One property, the dianionic headgroup of TOCL (69, 71, 72), could prevent SMA from binding to the membrane surface through electrostatic repulsion due to the negative charge density of SMA imparted by its maleic acid groups (104). The results of our SMA solubilization tests, however, suggest that charge repulsion is not a dominant factor in TOCL inhibition, at least under our experimental conditions. This is because: (i) LUVs containing the monoanionic POPG up to 30 mol% (with zeta potentials comparable to LUVs with 10-20 mol% TOCL) showed no inhibition of SMALP formation, and more compellingly, (ii) LUVs containing the dianionic MLCL (with zeta potentials equivalent to the cognate TOCL series) showed no SMA inhibition either. The second property, the inverse conical (H_II_-promoting) molecular geometry of TOCL (105), increases the lateral packing pressure in the acyl chain region (106, 107), which is known to inhibit SMA activity by hindering the insertion of its nonpolar styrene pendant groups into the nonpolar core of the membrane (32). Based on laurdan polarization in our LUV systems, TOCL increased lateral packing near the polar-apolar boundary, whereas MLCL trended toward the opposite effect, potentially explaining the inhibitory effect that TOCL had but MLCL did not. While the other test lipid POPE does have H_II_ propensity (108) and caused increased lipid packing based on laurdan polarization, it did not inhibit SMA activity in our system. The difference in SMA activity between these two LUV systems containing H_II_ lipids arise because TOCL has all unsaturated acyls, compared with POPE which is a mixed acyl (saturated/unsaturated) species, rendering more steric repulsion against SMA insertion in TOCL-containing LUVs in a way that is not reflected in the laurdan fluorescence profiles.

The marked effects of TOCL on SMALP formation that we observed in our system motivated us to further explore the physical basis of TOCL on SMA inhibition and how this lipid may affect the kinetics of SMA-dependent membrane disruption.

### 3.4. Early-stage kinetics of LUV solubilization by SMA

To address how TOCL may affect the initial stage of SMA-mediated membrane disruption, we devised a strategy for measuring solubilization in real time by pre-loading LUVs with the soluble pH sensitive dye 8-hydroxypyrene-1,3,6-trisulfonic acid (HPTS) (67, 109). This fluorescent probe reports pH based on an excitation wavelength ratio (λ_ex_ 405 nm / λ_ex_ 458 nm) at a constant emission wavelength (λ_em_ 509 nm), which can be calibrated against known pH standards (**Supplementary Figure S2**). By this approach, setting up a reaction with a pH differential between the medium and the vesicle lumen provides a readout of membrane integrity, as SMA disruption exposes the luminal probe to a different pH. Thus, we prepared LUVs of different TOCL content (up to 30 mol% TOCL) containing HPTS in the lumen at pH 7.3, then diluted the vesicles in an alkaline (pH 8.5) buffer to perform time course experiments (**Figure 5**). Before SMA addition, HPTS-loaded LUVs of all lipid compositions were near the vesicle preparation pH (∼7.3) and statistically identical, confirming that the LUV preparations were intact and proton-impermeant. Following SMA addition, samples containing POPC-only LUVs (blue trace) showed an immediate exponential rise in the pH reported by HPTS, suggesting that vesicles were efficiently disrupted with no detectable kinetic barrier, consistent with SMA time courses previously reported (32). By contrast, the LUVs containing 10 mol% TOCL (green trace) showed much slower disruption kinetics, and those with 20-30 mol% TOCL (orange and red traces) exhibited minimal or no SMA-induced leakage over the time course. Note that because this assay was meant to only address the initial kinetics of SMALP formation, the reactions did not likely reach a true end point, explaining why the HPTS signals of the POPC-only control samples did not stabilize or reach the pH 8.5 point prior to termination. These data suggest that TOCL impedes the initial steps of membrane permeabilization, likely by increasing lateral packing and bilayer mechanical stability.

**Figure 5.**
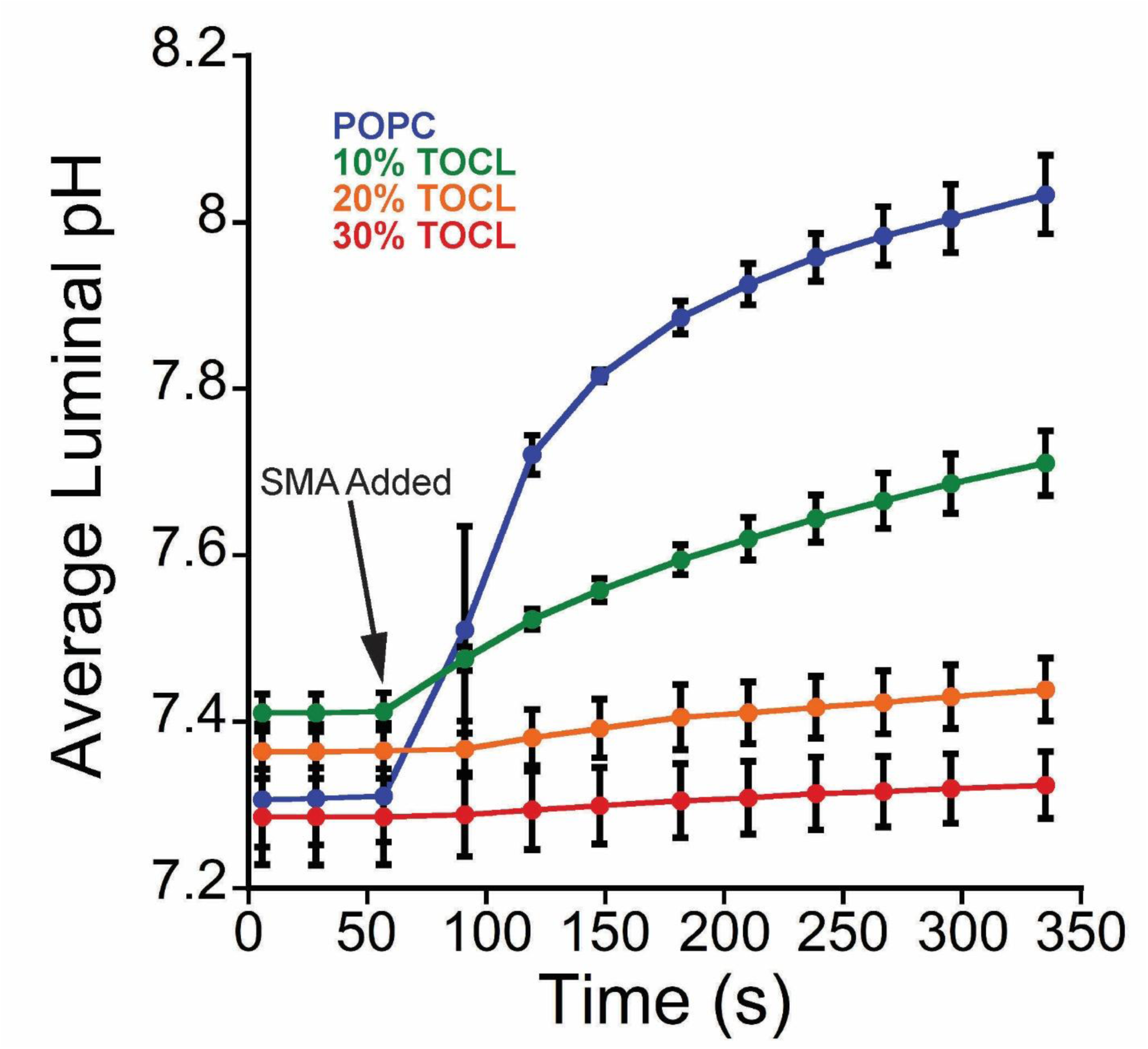
Kinetics of SMA-mediated disruption of TOCL-containing vesicles. LUVs composed of POPC only (blue), 10 mol% TOCL (green), 20 mol% TOCL (orange), and 30 mol% (red) containing luminal HPTS (pH 7.3) were incubated in reaction medium (pH 8.5) in a thermostatted fluorescence cuvette, with excitation readings made at 50 s intervals. SMA (final concentration 0.34 mg/ml) was added to the reaction at t = 56 sec (arrowhead). Individual data points represent means ± SD (n = 3-5 experimental replicates) and colored lines represent the reported pH change over time.

When considered alongside our DLS results (**Figure 4**), these findings suggest a model in which SMA-mediated solubilization is a multistep process, beginning with polymer binding and insertion, followed by progressive membrane deformation and nanodisc formation. The fact that nanodisc formation begins at similar SMA concentrations (**Figure 4C, inset**) despite suppressed leakage (**Figure 5**) suggests that early superficial interactions of SMA with the bilayer may occur without complete vesicle disruption. However, TOCL-enriched bilayers appear to resist both the early destabilization needed for leakage and the later membrane remodeling required for full solubilization. Together, these findings are consistent with the cooperative nature of bilayer solubilization by SMA; while preliminary steps can proceed unhindered, the inhibitory nature of TOCL-enriched bilayers can exert influence on the later stages of nanodisc formation.

### 3.5. Potentiation of SMALP formation using a bilayer-active alcohol

We next tested whether modulating the fluidity and lateral packing properties of TOCL-containing LUVs would make them less refractory to SMA solubilization. To this end, we incubated TOCL-containing LUVs with 1-heptanol prior to incubation with 3:1 SMA. Aliphatic alcohols intercalate between lipids and alter membrane physical properties by reducing packing pressure, enhancing elasticity, promoting disorder, and increasing area per lipid (decreasing bilayer thickness) (110–113). When pre-incubated with 5 mM 1-heptanol, LUVs containing 20 mol% TOCL became significantly more amenable to SMALP formation based on DLS analysis (**Figure 6A**). We observed that LUVs treated with 1-heptanol had an increased DLS-detected hydrodynamic radius, a known effect of alcohols on liposomes (114, 115) that may also cause the apparent increase in SMALP size. We addressed the underlying effects of 1-heptanol treatment on the physical properties of our LUVs, finding that this short chain alcohol reduced the laurdan-detected GP value (**Figure 6B**) and did not significantly change the surface charge detected by ζ readings (**Figure 6C**). Taken together, these results indicate that 1-heptanol treatment helped overcome the CL-imposed barrier to SMALP formation by reducing lateral lipid packing pressure and not by reducing SMA-membrane charge repulsion.

**Figure 6.**
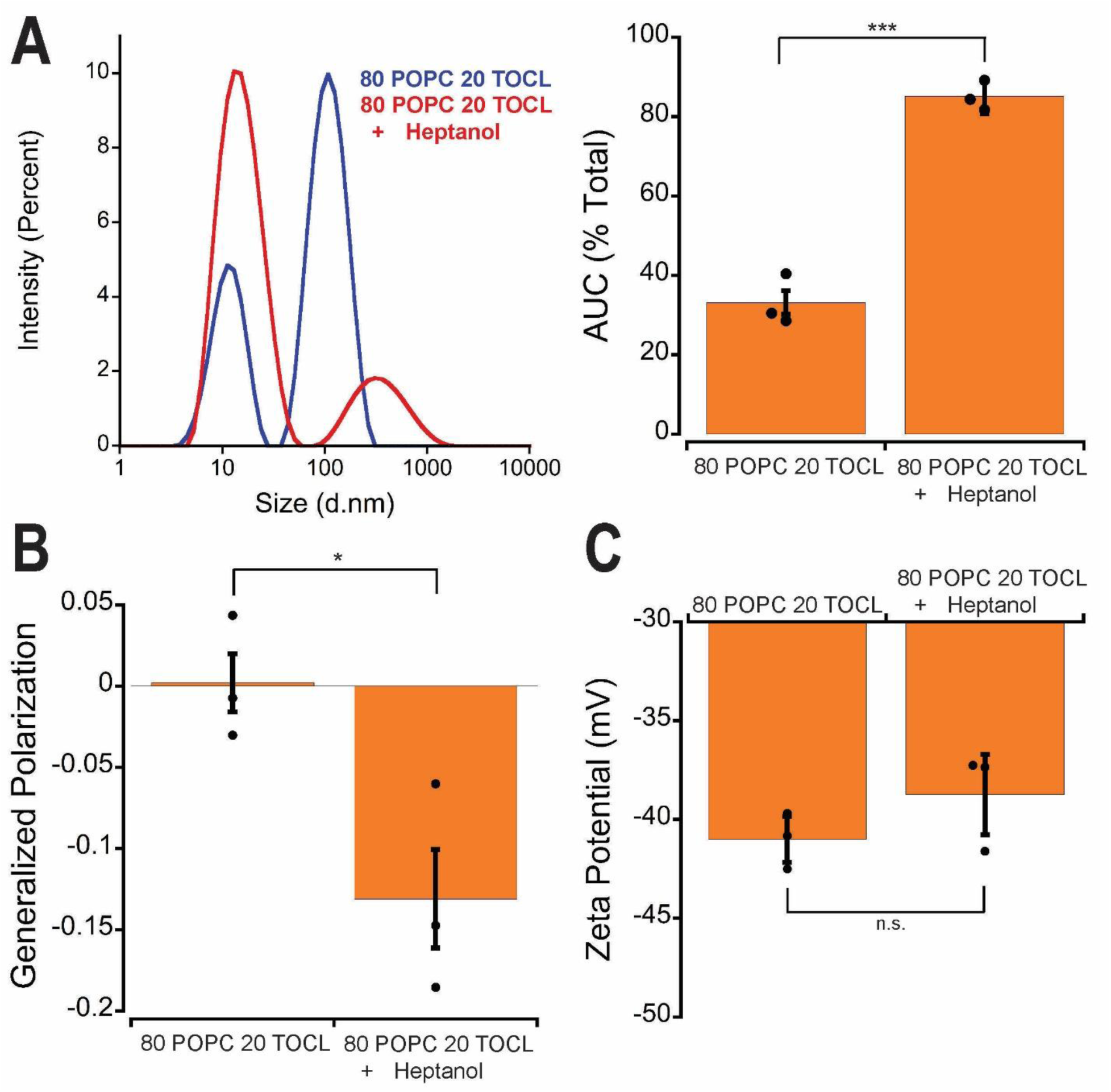
Effects of 1-heptanol on SMA solubilization of TOCL-containing vesicles. LUVs composed of 20 mol% TOCL were pre-incubated with or without 5 mM 1-heptanaol. For all measurements, individual data points represent means ± SD (n = 3-5 experimental replicates). (A) *Left,* intensity-weighted DLS size distribution of LUVs without (blue trace) or with (red trace) 1-heptanol. *Right*, quantitation of AUC corresponding to SMALPs as a fraction of total area from all particles by DLS size distribution measurements. Statistical significance was determined with an unpaired t-test (***P < 0.0005). (B) Laurdan-detected GP values determined as described in Figure 1. Statistical significance was determined with a paired t-test (*P < 0.05). (C) ζ readings determined as described in Table 1.

## 4. Conclusions

In this study, we have shown that among the four test phospholipids incorporated into POPC-based bilayers (POPG, POPE, TOCL, and MLCL), only TOCL substantially inhibited SMA-mediated membrane solubilization. Our data point to lateral acyl chain packing pressure, rather than charge-based repulsion, as the principal mechanism of inhibition. This conclusion is supported by three main lines of evidence: (i) LUVs containing other negatively charged lipids (POPG and MLCL) showed no such inhibition despite similar or greater anionic surface charge (verified by ζ measurements); (ii) fluorescence-based assessments of membrane order indicated that TOCL, but not its triacyl counterpart MLCL, markedly increased lipid packing near the polar-apolar boundary; and (iii) partial rescue of solubilization by the bilayer-active alcohol 1-haptanol further supports the interpretation that TOCL-induced ordering is a biophysical barrier to SMA insertion.

These findings expand our understanding of how lipid geometry and acyl chain properties influence SMA-membrane interactions. While charge repulsion has been well documented as a cause of SMA resistance, our results underscore the importance of lipid packing pressure, particularly in the context of non-bilayer-forming lipids like CL. Given the mitochondrial enrichment of CL, particularly in the inner membrane, these results may help explain the relative inefficiency of SMA-based extraction methods for mitochondrial proteins and highlight the need for novel copolymers tailored to overcome the physical barriers presented by non-lamellar lipids. Additionally, our results suggest that local membrane modifiers such as bilayer-active alcohols, might be exploited to potentiate SMA access to otherwise refractory membrane regions.

## CRediT authorship contribution statement

**Joseph Iovine:** Conceptualization, Data curation, Formal analysis, Investigation, Writing – original draft, Writing – review and editing.

**Nathan Alder:** Conceptualization, Formal analysis, Investigation, Project Administration, Funding acquisition, Writing – review and editing.

## Declaration of competing interest

The authors declare that they have no known competing financial interests or personal relationships that could have appeared to influence the work reported in this paper.

## Acknowledgements

This study was supported by the National Institutes of Health grant R01GM136975. We would like to thank. Dr. Steven M. Claypool and the members of the Alder Laboratory for fruitful discussions during the design, execution, and interpretation of results during this study.

---

**Supplementary Table 1.**

| **Variable Lipid** | **LUV Composition (% POPC/% Variable Lipid)** | **Polydispersity Index (Representative)** |
| --- | --- | --- |
| TOCL | 90/10 | 0.1741 |
|  | 80/20 | 0.1807 |
|  | 70/30 | 0.1814 |
| POPG | 90/10 | 0.1059 |
|  | 80/20 | 0.1230 |
|  | 70/30 | 0.1024 |
| POPE | 90/10 | 0.1224 |
|  | 80/20 | 0.1317 |
|  | 70/30 | 0.1619 |
| MLCL | 90/10 | 0.1176 |
|  | 80/20 | 0.1008 |
|  | 70/30 | 0.1175 |

**Supplementary Figure S1.**
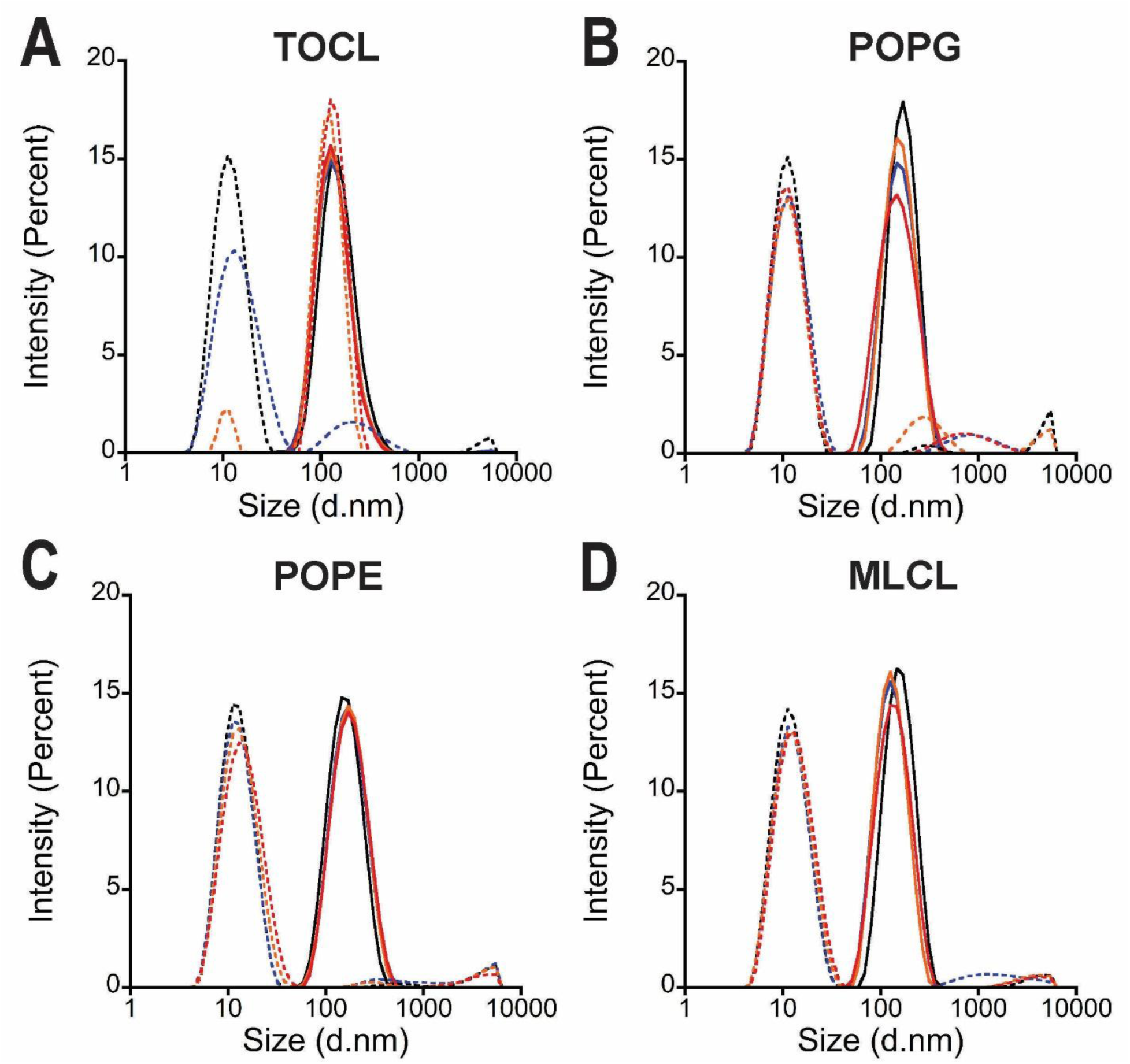
Representative intensity-weighted DLS size measurements of LUVs containing (A) TOCL, (B) POPG, (C) POPE, and (D) MLCL test lipids as indicated, before (solid lines) and following (dashed lines) SMA incubation, with test lipids at 0 mol% (black), 10 mol% (blue), 20 mol% (orange), and 30 mol% (red).

**Supplementary Figure S2.**
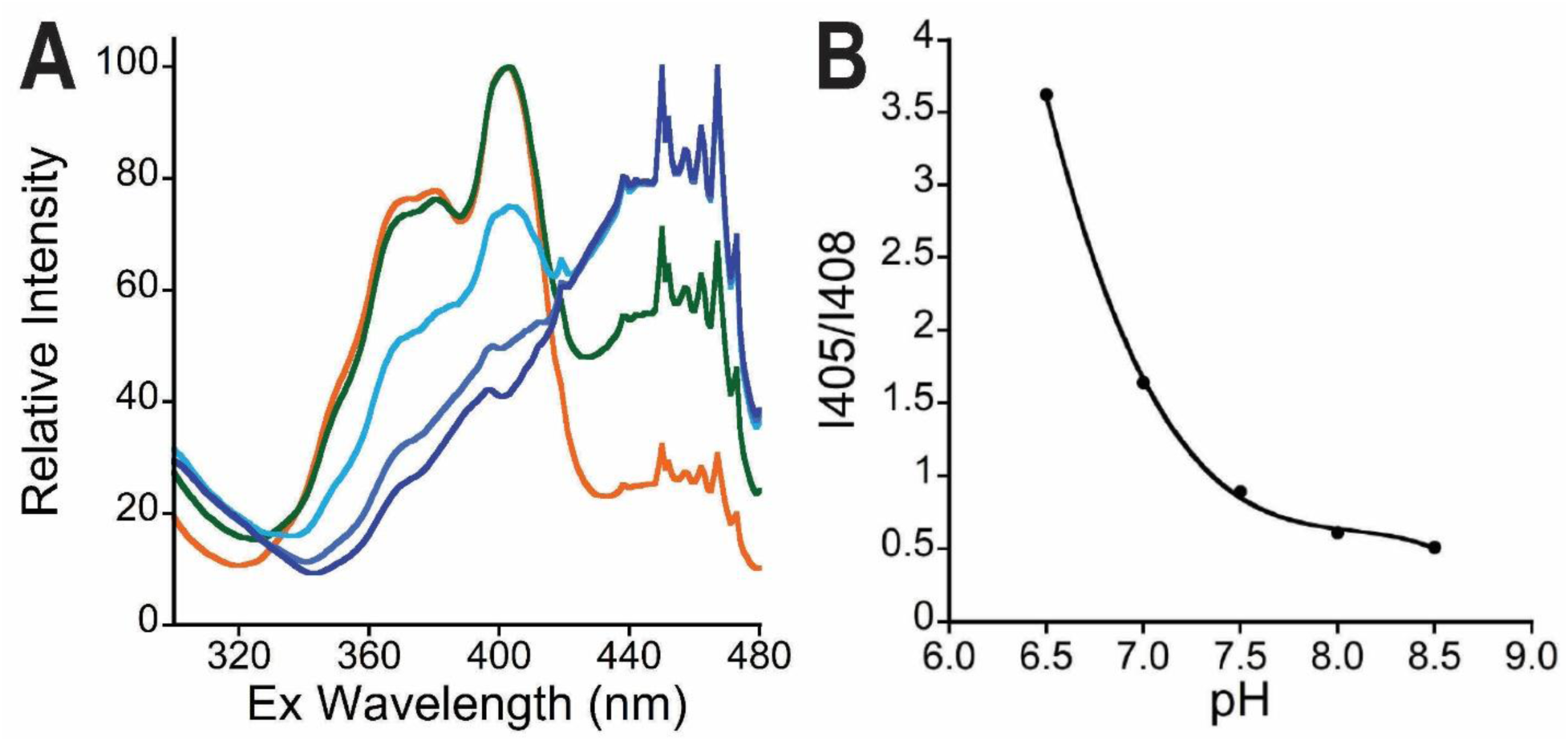
Calibration of pH readouts based on HPTS fluorescence. (A) Excitation scans (λ_ex_ X to X nm) at a fixed emission wavelength (λ_em_ 509 nm) using HPTS-loaded vesicles set to the experimentally verified solution pH values indicated. (B) Intensity ratios (λ_ex_ 405 nm / λ_ex_ 458 nm) at different pH values with points taken from scans in (A) and fit to the function indicated.

## Notes

### Competing Interest Statement

Nathan N. Alder reports financial support was provided by National Institutes of Health. If there are
other authors, they declare that they have no known competing financial interests or personal relationships that could have appeared to influence the work reported in this paper

